# Cotton growth, yield, quality and boron distribution as affected by soil-applied boron in calcareous saline soil

**DOI:** 10.1101/2020.04.02.021600

**Authors:** Atique-ur-Rehman, Rafi Qamar, Abid Hussain, Hassan Sardar, Naeem Sarwar, Hafiz Muhammad Rashad Javeed

**Affiliations:** Department of Agronomy, Bahauddin Zakariya University, Multan, Punjab, Pakistan; Department of Agronomy, College of Agriculture, University of Sargodha, Sargodha, Punjab, Pakistan; Department of Horticulture, Bahauddin Zakariya University, Multan, Punjab, Pakistan; Department of Environmental Sciences, COMSATS University Islamabad, Vehari Campus, Vehari 61100, Pakistan

**Keywords:** Cotton, soil application, photosynthesis, fiber quality, boron

## Abstract

Boron (B) is deficient in the calcareous, Typic Haplocambid soils of cotton growing belt of Pakistan, and thus is a vital reason for less cotton yield in the region. In order to investigate the growth and quality alterations associated with soil applied B on cotton (*cv. CIM-616 and CIM-600*) an experiment was conducted. Boron was applied at 0.00, 2.60, 5.52, 7.78 and 10.04 mg B kg^−1^ of soil using borax (Na_2_B_4_O_7_.10H_2_O), in a complete randomized design with factorial arrangement with four replications. Results revealed that soil applied B @ 2.60 mg B kg^−1^ of soil significantly (P≤0.05) improved cotton growth, yield, quality and B distribution among different parts. Different growth and yield parameters like plant height, leaf area, number of bolls, boll size and weight, seed cotton yield, photosynthesis, transpiration rate, stomatal conductance, water use efficiency, GOT, staple length and fiber fineness and strength except B uptake by roots, seed, leaves and stalk plant body which was significantly increased with B (10.04 mg B kg^−1^) in both cultivars of cotton, but the degree of effects was varied between cultivars. The results indicated that studied traits of both cultivars were significantly (P≤0.05) decreased in B-deficient stressed treatments. Between hybrids, CIM-600 produced significantly (P≤0.05) maximum recorded parameters under 2.60 mg B kg^−1^ application compared than CIM-616. Our findings confirm that the adequate level of B (2.60 mg B kg^−1^) had pronounced effects on various growth, yield, physiological and fiber quality associated traits, as compared to B uptake traits of cotton cultivars.

## Introduction

Pakistan ranked 4^th^ in share of production and consumption and 3^rd^ in export of cotton (*Gossypium hirsutium* L.) in the world [1]. However, yield stagnation (752 kg ha^−1^) and poor fiber quality are serious problems in cotton production [2]. Certain factors are particular for these problems, include poor fertilization especially lack of certain micronutrients like boron (B), which could improve the cotton yield and quality [3]; [4]. Boron deficiency has been prominent in cotton growing regions of the world including Pakistan, where 50% cotton growing area is deficient in B [5]; [6] and [7]. Its deficiency is common in tropical soils, where organic matter and clay content are lower [8] which are responsible for its leach down through the soil profile [9]; [10]. Degree of B adsorption onto the soil surfaces depend on the soil characteristics such as structure, pH, organic matter and clay content, iron and aluminum oxide and hydroxyl content and salinity [11]; [12]. However, its balanced application needs more consideration due to its narrow range between deficiency and toxicity which significantly retard the cotton production and physiological traits without any visible symptoms [13]. Moreover, B use improvement is difficult due to its low mobility in phloem vessels and result in low degree of its reutilization in cotton plant [14]; [15]. Due to low mobility in phloem vessels the required concentration of photosynthate and carbohydrate are not reached from leaves to fruits [15] that increases the rate of squares and bolls shedding at maturity, which finally affect the fiber quality [16]. Boron temporary deficiency can cause irreversible damage in cotton plants and thus significantly affect cotton yield [17]. Moreover, B withdrawal for a short period, establish deficiency and disturb the reproductive structures [18]; [19]. Likewise, boll retention depends on carbohydrates concentration in the plant body that mainly influenced by photoassimilate translocation from leaves to fruits, which under B deficiency decreases with increase in abortions [6]. Boron deficiency indirectly affects the metabolism of proteins and nucleic acids [20]; [21], and also mediates the levels of hormones and phenolic substances in the plant body [22]; [23] and [24]. Boron low mobility may cause temporary deficiency in cotton, although excess amount of B present in the soil solution. Due to critical role of B and lower mobility in cotton, continuous fertilization of B is needed throughout the plant’s life.

Boron fertilization not only improved the establishment and progress of reproductive organs [25]; [26] but also plays a crucial role in the vegetative growth of cotton plants [27]; [28]. However, continuous application of B without any soil test can also generate the toxicity in the soil [29], which disturb various physiological processes in cotton plants, such as reduction of chlorophyll contents, photosynthetic rates, lower cell division in root portion and lignin contents [30]. Boron uptake and transport in the new developing tissues depends on the transpiration stream, which may be reduced due to low evaporation rate, stomatal conductance in tropical region. However, there are some contradictory findings about B mobility within cotton plants, with respect to the most suitable rates and application forms [31]. A critical level of B concentrations in matured cotton leaf is 53 mg B kg^−1^, which should be 15-20 mg B kg^−1^ [4]. Due to narrow range of B concentration, plant analysis is not reliable technique for estimating of B nutritional status. Furthermore, new cotton genotypes response to B generally vary, although no significant differences were recorded among old cotton cultivars [32]; [31]. The variation in their abilities of carbohydrate transport, use and storage of B related mechanisms might be the reason [14]. Boron demands in cotton plants are relatively high compared to other crops [6] and it requires an average of 340 g B ha^−1^ from which about 12% is to be accumulated in seed [33]. Therefore, its slight excess may deteriorate the fiber quality [34]. In cotton growing regions, it is need of the time to find solution for B deficiency especially for the newly developed genotypes. The present study is therefore, conducted for improvement of yield and quality of cotton fiber in hyperthermic, sodic haplocambids, haplic Yermosols of cotton belt of Pakistan.

## Materials and methods

The experiment was conducted in earthen pots placed in wire-house at Bahauddin Zakariya University, Multan (30.10 °N, 71.25 °E and 421 ft. altitude above sea level) during cotton growing season 2018. Earthen pots (25 x 40 cm^−2^) filled with 20 kg soil and covered with polyethylene sheet having bulk density ≈ 1.04 mg m^−3^ [35]. Soil in the pots was equilibrated before 7 days of sowing [36]. Before conducting experiment, soil was air-dried, crushed and pass through 2 mm sieve for performing different physico-chemical properties. Hydrometer techniques was used for determination of soil textural class and it was silty clay loam belongs to Sindhalianwali soil series, and was hyperthermic, sodic haplocambids/Haplic Yermosols according to USDA and FAO classification, respectively. Soil pH and EC were 8.3 and 12 dS m^−1^ that were measured by a pH meter (Beckman 45 Modal, US) and EC meter (VWR Conductivity Meter DIG2052) respectively. Soil organic matter content was 0.78%, while total N 0.035%, NaHCO_3_-DTPA available-P 7.65 mg kg^−1^ and NH_4_OAc-extractable- K 162 mg kg^−1^. From soil analysis it is clear that B was deficient (0.43 mg B kg^−1^). Summary of weather data during the crop growth period is depicted in Fig. 1.

**Fig. 1.**
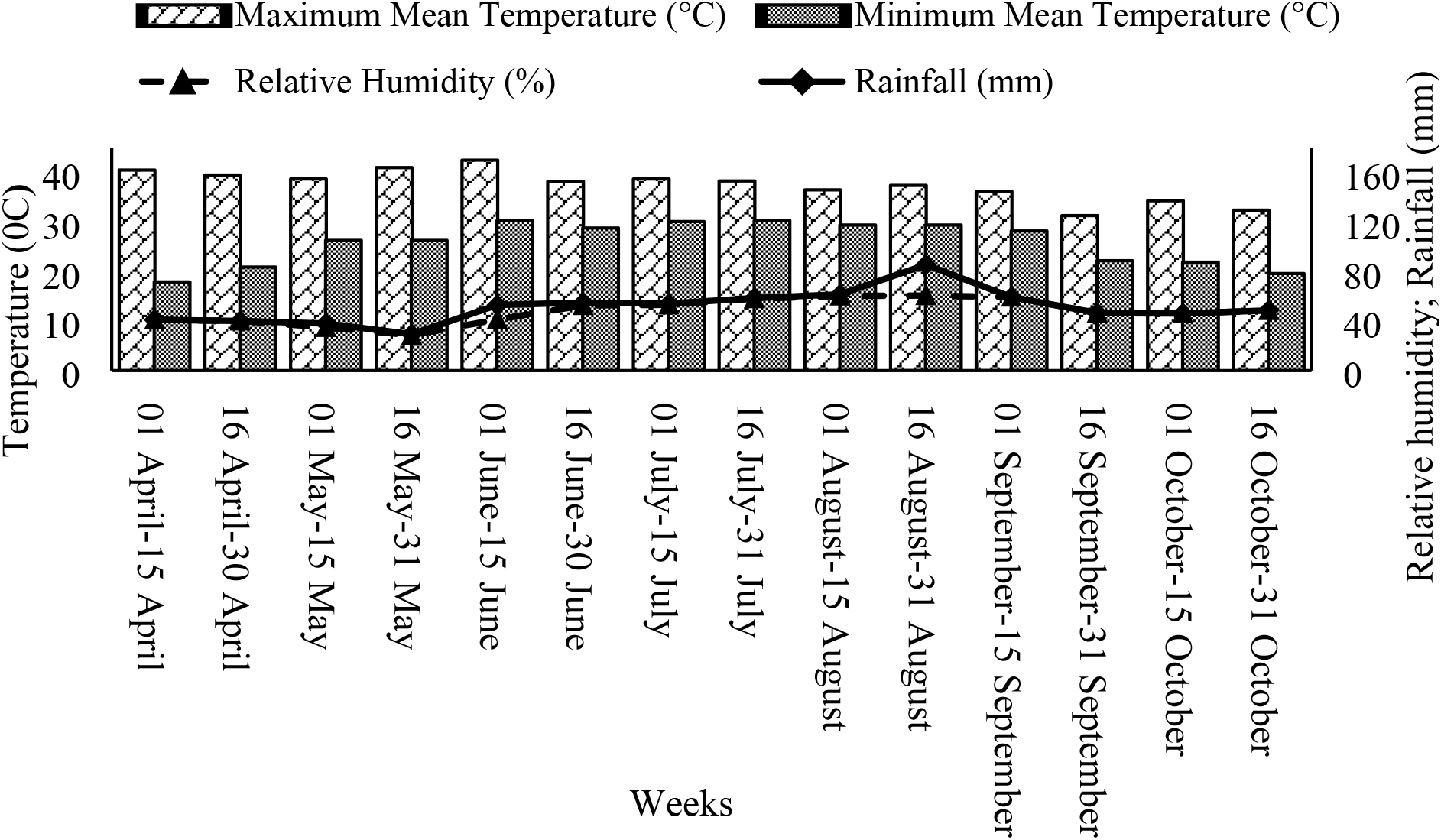
Meteorological data recorded at Bahauddin Zakariya University, Multan during 2018

The experiment was laid out according to completely randomized design (CRD) with factorial arrangement. There were two cotton cultivars viz. CIM-616 and CIM-600 tested with different B treatments viz. control or 0, 2.6, 5.52, 7.78 and 10.04 mg B kg^−1^ soil and each treatment was replicated five times. Boric acid (H_3_BO_3_) (17.5% B) was used as source of B. A total of 10 delinted cotton seeds of both cultivars were sown in each pot on 1^st^ May, 2018 and plants were thinned to two at 15 days after sowing (DAS). Soil in the pots were monitored and maintained at soil moisture up to 70% by weighing regularly. Recommended doses of N, P and K @ 200, 100 and 70 kg ha^−1^ were uniformly mixed thoroughly into the soil. When plants reached at maturity their bolls were separated and lint was detached manually from seed. Standard production practices were adopted and plants were kept free of insect-pests through using pesticide sprays.

For measuring different parameters, standard procedures were adopted. Leaf area of selected plants was determined with a leaf area meter (CI-202. Portable Laser Leaf Area Meter). Measurements of photosynthesis, stomatal conductance and transpiration rate of fully expanded leaves were taken between 9h00 and 11h30 with a portable photosynthesis measuring system (IRGA - LI-6400, LICOR). Water use efficiency was determined by dividing photosynthesis with transpiration rate. The plants were uprooted carefully at maturity and divided into roots, shoots, leaves and seed after removing lint. For determination of B concentration, different plant parts were separately washed with de-ionized water then dried in a thermo-ventilated oven at 65 ± 5°C up to constant weight. Then dried material was ground in a John Wiley mill and passed through a 40 mesh screen. The ground material was dry ashed at 550°C for 6 hours in a muffle furnace. Then the ash was taken in 0.36N H_2_SO_4_ and the B concentration was determined by spectrophotometer at 420 nm wavelength using azomethine-H method [37]. The samples of seed-cotton were separated into lint using single roller laboratory gin and then ginning out turn (GOT) was calculated. Fiber quality traits viz. fiber length, fiber fineness and fiber strength were analyzed on High Volume Instrument (HVI), manufactured by M/S Zellwegar Uster Ltd., Switzerland. The instrument was calibrated as per the instruction manual [38] followed by the standard procedure as described by [39].

Data collected were statistically analyzed by Fisher’s analysis of variance and treatment means were compared using least significant difference at 5% probability level [40].

## Results

Boron application to cotton had significant effects on cotton growth, yield and fiber quality (Table 1 to 4). However, plant height was affected by application of B and among different treatments, 2.6 mg B produced taller plants than other treatments (Table 1). From both cultivars, CIM-600 expressed 1% more plant height than CIM-616 (Table 1). Likewise, application of 2.6 mg B expressed higher leaf area than other treatments. Different yield contributing parameters were also improved more by 2.6 mg B application and an increase of 38.5% in number of bolls per plant was recorded with this treatment. Likewise, number of bolls per plant of CIM-600 was recorded 13% higher than CIM-616 (Table 1). An increase of 46% in boll size was also recorded with 2.6 mg B and it was remained well than all other treatments. The cultivar CIM-600 showed 16% bigger boll size than CIM-616 under different doses of soil applied B (Table 1). Soil applied 2.6 mg B in CIM-600 and CIM-616 produced 52% heavier boll weight CIM-600 and CIM-616 (Table 1). Similarly, maximum seed cotton yield per plant was recorded by 2.6 mg B that was 51% more than control. Between two cultivars, a 2% higher seed cotton yield per plant was recorded in CIM-600 than CIM-616 (Table 1).

**Table 1:** Influence of soil applied boron on number of bolls per plant, boll size and weight and seed cotton yield per plant of cotton cultivars

| Treatments<br>(mg B kg<sup>-1</sup><br>soil) | Plant height (cm) |  |  | Leaf area (cm) |  |  | Number of bolls per<br>plant |  |  | Boll size (cm) |  |  | Boll weight (g) |  |  | Seed cotton per plant (g) |  |  |
| --- | --- | --- | --- | --- | --- | --- | --- | --- | --- | --- | --- | --- | --- | --- | --- | --- | --- | --- |
|  | CIM-616 | CIM-600 | Mean | CIM-616 | CIM-600 | Mean | CIM-616 | CIM-600 | Mean | CIM-616 | CIM-600 | Mean | CIM-616 | CIM-600 | Mean | CIM-616 | CIM-600 | Mean |
| 0.0 | 86.8<br>d | 87.7c | 87.3B | 143.6<br>f | 144.2<br>ef | 143.9<br>C | 15i | 18g | 16E | 1.8f | 2.3de<br>f | 2.1D | 2.07e | 2.83c<br>d | 2.45D | 220.25g | 225.54g | 222.89<br>E |
| 2.6 | 90.7<br>b | 91.3a | 91.0A | 146.5<br>b | 147.6<br>a | 147.1<br>A | 24c | 29a | 26A | 3.4b | 4.4a | 3.9A | 4.30a | 4.50a | 4.40A | 448.90b | 460.52a | 454.71<br>A |
| 5.52 | 86.4<br>d | 88.1c | 87.3B | 144.5<br>de | 145.6<br>c | 145.0<br>B | 23d | 26b | 24B | 3.1bc | 3.5b | 3.3B | 3.20b<br>c | 3.50b | 3.35B | 418.19c | 426.92c | 422.55<br>B |
| 7.78 | 83.3<br>f | 84.2e | 83.7C | 143.5<br>f | 145.2<br>cd | 144.3<br>C | 21e | 24c | 22C | 2.6cd<br>e | 2.8cd | 2.7C | 2.73d | 3.16b<br>c | 2.95C | 308.12e | 322.45d | 315.28<br>C |
| 10.04 | 80.4<br>h | 81.3g | 80.9D | 143.7<br>f | 144.6<br>de | 144.2<br>C | 17h | 21e | 19D | 2.1ef | 2.7cd | 2.4C<br>D | 2.16e | 2.30e | 2.23D | 238.42f | 245.58f | 242.00<br>D |
| Mean | 85.5<br>B | 86.5A |  | 144.4<br>B | 145.4<br>A |  | 20B | 23A |  | 2.6B | 3.1A |  | 2.89B | 3.26A |  | 327.16<br>B | 332.82<br>A |  |
| LSD at 5<br>% | Cultivars (C):<br>0.086; Boron levels<br>(B): 0.28; C x B:<br>0.47 |  |  | Cultivars (C): 0.24;<br>Boron levels (B):<br>0.46; C x B: 0.77 |  |  | Cultivars (C): 0.14;<br>Boron levels (B):<br>0.35; C x B: 0.59 |  |  | Cultivars (C): 0.05;<br>Boron levels (B):<br>0.33; C x B: 0.56 |  |  | Cultivars (C): 0.05;<br>Boron levels (B):<br>0.26; C x B: 0.45 |  |  | Cultivars (C): 4.22; Boron<br>levels (B): 3.76; C x B:<br>6.33 |  |  |

Different physiological traits were significantly improved with B application and among different treatments, 2.6 mg B produced 45% higher photosynthesis than control, and was significantly higher than other B treatments. From both cultivars, CIM-600 produced 8.2% higher photosynthesis than CIM-616 (Table 2). Almost same trend was recorded for transpiration rate. Likewise, application of 2.6 mg B produced 9% more transpiration rate than control, which was significantly higher than other B treatments (Table 2). Between two cultivars, transpiration rate of CIM-600 was 0.50% more than CIM-616 (Table 2). Regarding stomatal conductance, 37% higher stomatal conductance than control was recorded application of 2.6 mg B, which was significantly higher than other treatments. From both cultivars, CIM-600 recorded 1% higher stomatal conductance than CIM-616 (Table 2). Similarly, application of 2.6 mg B improved water use efficiency, that was 40% higher than control and other treatments. From both cultivars, 9% higher water use efficiency was recorded in CIM-600 than CIM-616 (Table 2).

**Table 2:** Influence of soil applied boron on photosynthesis, transpiration, stomatal conductance and water use efficiency of cotton cultivars

| Treatments<br>(mg B kg<sup>-1</sup> soil) | Photosynthesis<br>($\mu\text{mol CO}_2 \text{ m}^{-2} \text{ s}^{-1}$) | | | Transpiration rate<br>($\text{mmol m}^{-2} \text{ s}^{-1}$) | | | Stomatal conductance<br>($\mu\text{mol m}^{-2} \text{ s}^{-1}$) | | | Water use efficiency<br>($\mu\text{mol CO}_2 \text{ mol}^{-1} \text{ H}_2\text{O day}^{-1} \text{ m}^{-2}$) | | |
| --- | --- | --- | --- | --- | --- | --- | --- | --- | --- | --- | --- | --- |
|  | CIM-616 | CIM-600 | Mean | CIM-616 | CIM-600 | Mean | CIM-616 | CIM-600 | Mean | CIM-616 | CIM-600 | Mean |
| 0.0 | 5.01j | 5.42i | 5.22E | 13.20h | 13.40g | 13.30E | 2.36e | 2.40e | 2.38E | 0.38h | 0.40g | 0.39E |
| 2.6 | 9.12b | 9.92a | 9.52A | 14.61b | 14.68a | 14.64A | 3.74a | 3.77a | 3.76A | 0.62b | 0.68a | 0.65A |
| 5.52 | 7.92d | 8.32c | 8.12B | 13.92d | 13.97c | 13.94B | 3.60b | 3.62b | 3.61B | 0.57d | 0.59c | 0.58B |
| 7.78 | 6.42f | 7.82e | 7.12C | 13.78e | 13.81e | 13.79C | 2.91c | 2.95c | 2.96C | 0.47e | 0.57d | 0.52C |
| 10.04 | 6.12h | 6.22g | 6.17D | 13.50f | 13.52f | 13.51D | 2.83d | 2.87d | 2.85D | 0.45f | 0.46ef | 0.45D |
| Mean | 6.92B | 7.54A |  | 13.80B | 13.87A |  | 3.09B | 3.12A |  | 0.49B | 0.54A |  |
| LSD at 5 % | Cultivars (C): 0.001; Boron levels (B): 0.007; C x B: 0.01 |  |  | Cultivars (C): 0.01; Boron levels (B): 0.17; C x B: 0.02 |  |  | Cultivars (C): 0.01; Boron levels (B): 0.03; C x B: 0.05 |  |  | Cultivars (C): 0.01; Boron levels (B): 0.01; C x B: 0.01 |  |  |

Boron contents in different parts of cotton was increased with increase in B application and there was an increasing trend of B in cotton roots, leaves, stalk and seed from lower to higher level (Table 3). For example, 74% higher B was recorded in roots with 10.04 mg B than control (Table 3). Similarly, 86% higher B contents were noted in leaves of CIM-600 with 10.04 mg B than other treatments (Table 3). The cultivar CIM-600 produced 2% higher B uptake by leaves than CIM-616 under different B treatments (Table 3). Similarly, maximum concentration of B in stalk was recorded at 10.04 mg B that was 67% higher than control (Table 3). Between two cultivars, a 4% higher B uptake by stalk was recorded in CIM-600 than CIM-616 (Table 3). Among different soil applied B, 10.04 mg had 40% higher B uptake by seed cotton than control (Table 3). The cultivar CIM-600 showed 3% higher B uptake by seed cotton than CIM-616 under different doses of soil applied B (Table 3).

**Table 3:** Influence of soil applied boron on boron uptake by roots, seed, leaves and stalk of cotton cultivars

| Treatments<br>(mg B kg<sup>-1</sup> soil) | Boron uptake by roots<br>(mg kg<sup>-1</sup>) |  |  | Boron uptake by seed cotton<br>(mg kg<sup>-1</sup>) |  |  | Boron uptake by leaves<br>(mg kg<sup>-1</sup>) |  |  | Boron uptake by stalk<br>(mg kg<sup>-1</sup>) |  |  |
| --- | --- | --- | --- | --- | --- | --- | --- | --- | --- | --- | --- | --- |
|  | CIM-616 | CIM-600 | Mean | CIM-616 | CIM-600 | Mean | CIM-616 | CIM-600 | Mean | CIM-616 | CIM-600 | Mean |
| 0.0 | 10.75j | 12.08i | 11.41E | 1.12j | 1.16i | 1.14E | 37.78j | 38.12i | 37.95E | 21.85j | 24.20i | 23.02E |
| 2.6 | 28.72h | 30.25g | 29.48D | 1.32h | 1.35g | 1.34D | 93.81h | 95.45g | 94.63D | 39.58h | 41.12g | 40.35D |
| 5.52 | 35.74f | 37.12e | 36.43C | 1.43f | 1.48e | 1.46C | 192.81f | 196.87e | 194.84C | 58.34f | 61.12e | 59.73C |
| 7.78 | 39.67d | 41.82c | 40.74B | 1.65d | 1.70c | 1.68B | 212.78d | 222.42c | 217.60B | 64.78d | 66.88c | 65.83B |
| 10.04 | 43.56b | 45.02a | 44.29A | 1.87b | 1.90a | 1.89A | 268.78b | 269.82a | 269.30A | 69.34b | 72.12a | 70.73A |
| Mean | 31.68B | 33.25A |  | 1.48B | 1.52A |  | 161.19B | 164.54A |  | 50.77B | 53.08A |  |
| LSD at 5 % | Cultivars (C): 0.01; Boron levels (B): 0.003; C x B: 0.005 |  |  | Cultivars (C): 0.002; Boron levels (B): 0.002; C x B: 0.003 |  |  | Cultivars (C): 0.001; Boron levels (B): 0.002; C x B: 0.003 |  |  | Cultivars (C): 0.03; Boron levels (B): 0.02; C x B: 0.04 |  |  |

Fiber quality traits were also significantly affected by B application (Table 4). Soil applied 2.6 mg B produced 6% higher GOT than control and other B levels. CIM-600 recorded 1.5% higher GOT than CIM-616 under different soil applied B (Table 4). Boron application significantly affected staple length and 2.6 mg B gave 3.5% more staple length than control (Table 4). From the cultivars, CIM-600 presented 5.4% higher staple length than CIM-616 (Table 4). Regarding fiber fineness, among different B concentrations, application of 2.6 mg and 5.52 mg B exhibited 17% higher staple length than control. The cultivar CIM-600 produced 15.5% higher fiber fineness than CIM-616 (Table 4). Likewise, 5% higher fiber strength was recorded with 2.6 mg B as compared to control while other B treatments were statistically at par with each other. Cotton cultivars CIM-600 attained 1.8% more fiber strength than CIM-616 (Table 4).

**Table 4:**
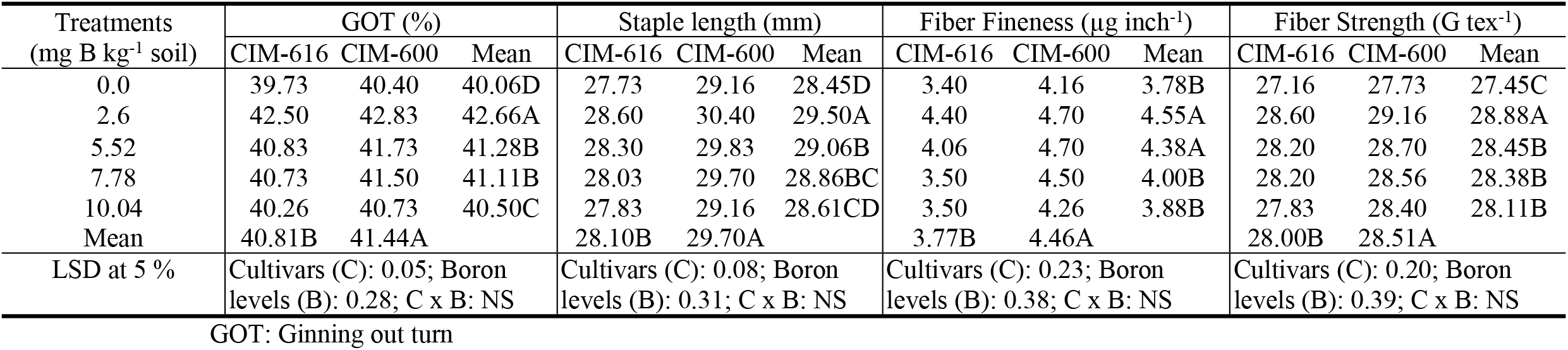
Influence of soil applied boron on GOT, staple length, fiber fineness and strength of cotton cultivars

## Discussion

Boron application has significant effects on growth, yield, physiological and fiber traits of cotton. Significant improvement in cotton plant height (11.1%) and leaf area (2.2%) with 2.6 mg B (Table 1) shows that appropriate dose of B paly role in different physiological, biochemical, metabolic and enzymatic activities of plant [41], thus its deficiency declines (4.1%) in plant height [42]. Moreover, lower plant height at higher B may be due to its narrow range between deficiency and toxicity which may damage the plant structure that limit the cotton growth without any visible symptoms [13]; [31]. Improvement in plant growth might be due to improved macronutrient uptake in response to B application [43]. Its deficiency severely declines various physiological and growth parameters like leaf area and seed cotton yield [42]. Boron insufficiency results in growth impairments such as reduced plant growth [6]. Higher number of bolls per plant and boll size and weight might be as result of increase in sugar translocation, membrane permeability, photosynthetic rate and migration of photosynthate from source to sink. [44] postulated that enhanced B supply to plants promotes flower development, pollen germination, fertilization, and seed development and, thus, reduces fruit shedding, which resulted in an increase in the number of bolls per plant and boll weight [45]; [19]. Boron deficiency causes significant shedding of square and boll shedding [4]; [46]. Seed cotton yield enhanced with the increase of boll size and weight, bolls per plant and B uptake (Table 3) and water use efficiency [6]; [47]; [4] and [46]. Abortion of reproductive parts occurred because of B deficiency that impairs the formation of the peduncle vascular system, impairing carbohydrate transport to the ovary [19]; [15]. Furthermore, higher concentrations of B have adverse effects on plant metabolic activities related to chlorosis and necrosis, loss of photosynthetic capacity and eventually reduction in plant productivity [48]. Necrotic areas developed on leaves between veins and loss of leaves due to toxicity, which inhibited photosynthetic process and exerted a negative impact on cotton growth [48].

Soil applied has significantly improved photosynthesis and other related parameters of cotton (Table 2). Earlier it is reported that photosynthesis improved 45.2% at 2.6 mg B/ Kg of soil in cotton (Table 2) which might be due to enhance in net assimilation rates which in turn is a measure of photosynthetic activity [49]. Significantly lower photosynthesis rate was possibly due to reduced chlorophyll biosynthesis as suggested by [46] in cotton. Our results supported the findings of [50] who reported that photosynthetic rate were lowered in plants under B deficiency leading to growth inhibition. The reduction in leaf area (Table 1) is mainly responsible for the lower photosynthetic rate in cotton plants under control conditions. Additionally, it has also been reported that B deficiency (control) reduces photosynthetic efficiency by changing stomatal density and stomatal conductance to decrease the conductivity of CO_2_ [51]. Higher rate (above 2.6 mg B) showed 35.2% lower photosynthesis, leaf chlorophyll contents, root cell division and lignin and suberin levels [30]. Present study showed 9.2%, 36.7% and 40% lower transpiration rate, stomatal conductance and water use efficiency in control than 2.6 mg B respectively. Our results validate the findings of [52] who concluded that significant decreased in stomatal conductance, photosynthetic rate and transpiration rate in the functional leaves of cotton after the formation of brown rings on the petiole are due to destruction of vascular bundle of petiole. [53] also reported that B deficiency deformate the phloem sieve which affects the transport of carbohydrates, water and nutrients that results in reduction of stomatal conductance and transpiration rate. Therefore, the destruction of petiole vessels impairs the transport of photosynthetic products of the leaves to the other parts. Significant reduction of transpiration rate and stomatal conductance in control also reduced the water use efficiency (WUE) (Table 2). Boron application raised WUE by up to 40% as compared to control (Table 2). Owing to other improvements in physiological performance, [51] reported that B enhanced stomatal conductance and reduced intercellular CO2 concentration and resulting a significant increase in photosynthesis, transpiration rate, stomatal conductance and WUE. Boron deficiency cause plant morphological changes, especially in the leaves [19] with a decrease in the number and functioning of stomata [54], which impairs transpiration rate [6].

Boron partitioning in different parts of cotton was varied among tissues. The B contents in roots, stem, leaves and seeds were increased with increase in B concentration in soil. It was noted that more B was in leaves than stalk that might be due to low mobility of the nutrient in the cotton phloem [19], owing to high transpiration rate, the main driving force of B transport within the plant [55]. The amount of B accumulated in the plant body increased exponentially with 2.6, 5.52, 7.78 and 10.04 mg B Kg^−1^ of soil (Table 3). Lower concentration of B in seeds as compared to other plants parts (Table 3) might be due to B movement through xylem, which has no direct connection to seed [56]. Moreover, flowers and seeds may not be able to take up B directly from the soil [57]; [58] and [45]. The partitioning of B in various plant tissues showed significant variations with increasing B level and it was assimilated in the order of leaf > shoot> root (Table 3) [59].

Different fiber quality traits of both cultivars were improved with soil applied B in calcareous soil (Table 4). Maximum GOT, staple length, fiber fineness and strength were recorded with 2.6 mg B (Table 4). The results were confirmed by the findings of [14] who reported that application of B enhanced the GOT, staple length and fiber fineness and strength of cotton genotypes but the degrees of effects were varied among genotypes. In present study, from the two cultivars, CIM-600 produced better quality fiber at different levels of B. Moreover, higher concentration (above 2.6 mg) of B deteriorates the fiber quality (Table 4). Our results are quite in line with the findings of [60], who reported that fiber quality was positively affected by application of B. Although fiber quality was improved with different treatments, however, it was the best with 2.6 mg B. Our results supported the findings of [61] who reported that soil applied B affected staple length, fiber fineness and strength and uniformity ratio.

## Conclusion

Boron application into the medium of cotton growing considerably improved the growth, different yield and quality parameters. Different gas exchange parameters were also improved, which ultimately improved the performance of cotton in saline soil. From the tow cotton genotypes, CIM-600 showed better results as compared to CIM-616. Among different B applications, 2.6 mg B Kg^−1^ of soil was remain superior to others.

## Acknowledgments

We acknowledge the financial support for this study from Directorate of Research, Bahauddin Zakariya University, Multan, Pakistan.

